# Genetic polymorphisms lead to major, locus-specific, variation in piRNA production in mouse

**DOI:** 10.1101/2022.10.21.513296

**Authors:** Eduard Casas, Pío Sierra, Cristina Moreta-Moraleda, Judith Cebria, Ilaria Panzeri, J. Andrew Pospisilik, Josep C. Jimenez-Chillaron, Sonia V. Forcales, Tanya Vavouri

## Abstract

PIWI-interacting RNAs (piRNAs) are small noncoding RNAs that silence transposons in the animal germline. PiRNAs are produced from long single-stranded non-coding transcripts, from protein-coding transcripts, as well as from transposons. While some sites that produce piRNAs are in deeply conserved syntenic regions, in general, piRNAs and piRNA-producing loci turnover faster than other functional parts of the genome. To learn about the sequence changes that contribute to the fast evolution of piRNAs, we set out to analyse piRNA expression between genetically different mice. Here we report the sequencing and analysis of small RNAs from the mouse male germline of four classical inbred strains, one inbred wild-derived strain and one outbred strain. We find that genetic differences between individuals underlie variation in piRNA expression. We report significant differences in piRNA production at loci with endogenous retrovirus insertions. Strain-specific piRNA-producing loci include protein-coding genes. Our findings provide evidence that transposable elements contribute to inter-individual differences in expression, and potentially to the fast evolution of piRNA-producing loci in mammals.

## Introduction

Eukaryotic genomes are hosts to a great number and diversity of transposons, that vary between species and between individuals of the same species (for a review see (Cosby et al., 2019). Active transposons are mobile genetic elements that can propagate within the genome of a cell. When a transposon replicates in the germline, the new copy is passed onto the next generation. To counter potentially deleterious, heritable, mutagenic events caused by transposons, living organisms have evolved mechanisms that repress these elements in the germline. One of the most important defence mechanisms protecting the animal germline against transposons is the PIWI-interacting RNA (piRNA) pathway (reviewed in (Ozata et al., 2019; Siomi et al., 2011)). The core components of the piRNA pathway are deeply conserved and active in the germline of almost all animals. Yet, even closely related mammalian species produce distinct sets of piRNAs. The genetic mechanisms that lead to the fast diversification and divergence of piRNAs between species are largely unknown.

The mammalian piRNA pathway surveys the germline for transposon transcripts and transposon-containing genomic loci using millions of piRNAs, each with distinct sequence. PiRNAs are small non-coding RNAs produced from a few hundred loci (Gainetdinov et al., 2018). Approximately half of these loci are non-coding, while the other half are protein-coding genes (Li et al., 2013). Why some germline-expressed coding or non-coding transcripts become processed into piRNAs and others do not remains unclear. A mechanism that defines a transcript as a piRNA precursor is the presence of sequence with extensive complementarity to initiator piRNAs produced from other genomic loci (Gainetdinov et al., 2018). Yet, this requires constant expression of piRNAs throughout the life cycle of the mammalian germline, which is not the case, suggesting that there are additional mechanisms that trigger piRNA production during development and during evolution.

The evolution of piRNAs is fast. Some piRNA-producing loci are found in syntenic regions of distantly related species, however their sequence is not conserved (Girard et al., 2006; Chirn et al., 2015a). Furthermore, the sites from which piRNAs are produced are more often than not, species-specific (Assis & Kondrashov, 2009; Chirn et al., 2015; Özata et al., 2020). Considering that approximately half of all mouse piRNA precursors are transcripts of protein-coding genes it is remarkable how evolvable production of piRNAs is.

Considering the fast evolution of piRNAs and piRNA-producing genes, how different are the sets of piRNAs expressed in genetically different individuals of the same species? There are few studies addressing this question, none of which in mammals. Analyses of piRNAs from different Drosophila (Kelleher & Barbash, 2013; Shpiz et al., 2014) and zebrafish strains (Kaaij et al., 2013) have revealed that the identity of piRNA-producing loci and their expression levels vary depending on their genetic background. In Drosophila, the comparison of two strains revealed that transposable element insertions in euchromatic regions induce the formation of dual-strand piRNA-producing loci at the site of insertion and single-strand piRNAs downstream of transposable element insertions in 3’ UTRs of protein-coding genes(Shpiz et al., 2014). Although the piRNA pathway is conserved between flies and mice, the mechanisms of piRNA biogenesis are quite distinct(ElMaghraby et al., 2019; Kneuss et al., 2019). Even though piRNAs have been extensively studied in mice and play an essential role in male fertility, it is unknown whether there are differences in the loci that produce piRNAs in different mice, or different individuals of any other mammalian species.

We sought to quantify the variation in piRNA expression in different strains of mice and then use it to search for potential genetic mechanisms for this variation. We sequenced and analysed small RNAs from the male germline of 57 adult mice from four classical inbred mouse strains (C57BL/6J, 129S1/SvImJ, C3HeB/FeJ, NOD), one wild-derived inbred strain (CAST/EiJ) and one outbred strain (ICR). We found significant differences in piRNA production from different genomic loci between genetically diverse mice and only minimal differences between mice of the same inbred strain. We tested the link between variation in piRNA expression and transposable element insertions or deletions and found a highly significant association, specifically for the murine endogenous retrovirus (ERV) Intracisternal A particle (IAP). Taken together with the previous work in fruitflies, our work in mice reveals that new transposable element insertions are a deeply conserved genetic mechanism for piRNA diversification within a species and the emergence of new piRNA-producing loci during evolution.

## Results

### Variation in piRNA expression between genetically diverse mice

We set out to analyse inter-strain variation in small RNA production from known piRNA-producing loci (also known as piRNA clusters) of the mouse genome, aiming to understand the level of piRNA expression variation between genetically diverse individuals in a mammalian species. As a first approach, we studied inter-individual variation in abundance of small RNAs mapped to 214 known piRNA producing loci (Li et al., 2013) from RNA extracted from whole testes of young adult mice from four classical inbred strains: C57BL/6J (referred to as BL6), NOD, C3HeB/FeJ (referred to as C3H) and 129S1/SvImJ (referred to as 129) (**Fig 1** and **Supplementary Tables 1-3**). The majority (75-80%) of small RNAs sequenced from whole testis from all four inbred strains mapped to known piRNA clusters (**Supplementary Table 1**). For brevity, we refer to small RNAs mapping to known piRNA clusters as piRNAs, even though these small RNAs were not identified bound to PIWI proteins by us. The reference set contains 84 loci producing piRNAs predominantly during the prepachytene stages of meiosis, 100 loci producing piRNAs predominantly from the pachytene stage of meiosis onwards and 30 loci that are known as hybrid, expressed throughout adult spermatogenesis (Li et al., 2013). As expected, the loci with the highest count of mouse piRNAs in the whole adult mouse testis in all four mouse strains are the pachytene ones (**Supplementary Fig 1**). Overall, piRNA abundance was highly correlated between animals of the four classical inbred strains (**Fig 1A,B**).

**Figure 1.**
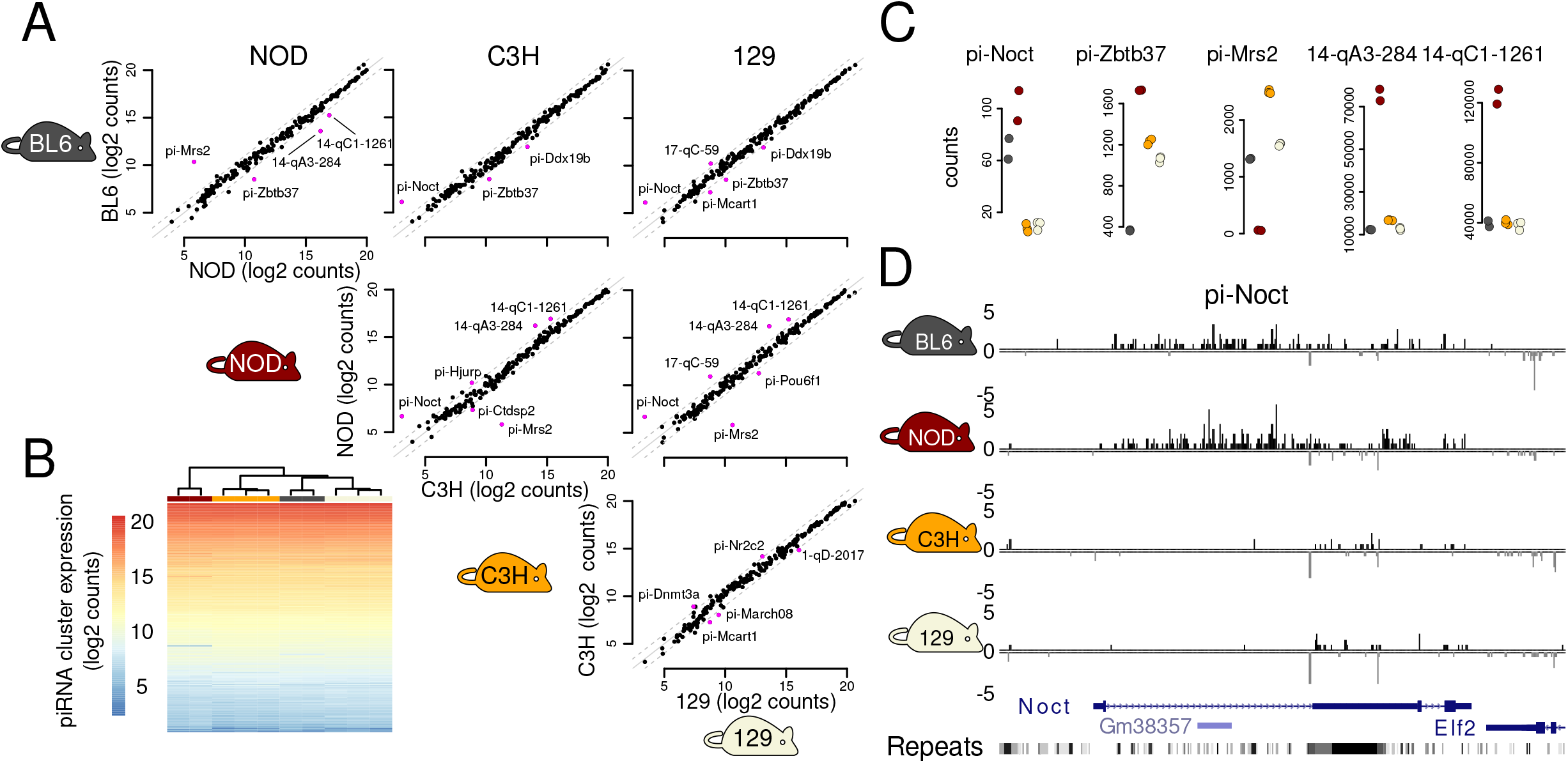
Variation in expression of 214 previously defined piRNA-producing loci in testes of four classical inbred mouse strains. (**A**) Pairwise comparison of expression of 214 previously defined piRNA-producing loci in four classical inbred mouse strains. Significantly differentially expressed piRNA clusters are shown using purple points and their names are shown. Samples from the same strain are averaged and expression values are scaled. (**B**) Heatmap of piRNA cluster expression in each mouse sample of the four inbred strains. (**C**) Five piRNA-producing loci with highly variable expression in four classical inbred mouse strains. Data points show the scaled small RNA counts with BL6 samples shown in grey, NOD in red, C3H in yellow and 129 in beige. Small RNAs mapped at these five loci on each strand and in each replicate are shown in Supplementary Figure 2. (**D**) Classical inbred mouse strains produce significantly different levels of small RNAs from pi-Noct (also known as pi-Ccrn4l). Genes and repeats are also shown. One sample from each strain was randomly chosen. The 214 piRNA-producing loci were defined in BL6 (Li et al., 2013).

Despite the high concordance of piRNA expression between all the studied mice, the abundance of piRNAs was more similar between animals of the same inbred strain than animals of different strains (**Fig 1B**). This observation prompted us to explore whether genetic differences between strains may explain variation in piRNA abundance from some clusters (“piRNA cluster expression”). Of the 214 known piRNA clusters, fifteen were significantly differentially expressed in at least one pairwise comparison and five of these clusters were significantly differentially expressed in at least three of the six pairwise comparisons between the four inbred mouse strains (**Fig 1A,C**); these are the clusters overlapping the protein-coding genes *Noct* (also previously known as *Ccrn4l*), *Zbtb37* and *Mrs2* and the noncoding clusters 14-qA3-284 and 14-qC1-1261, both on chromosome 14. The protein-coding gene *Nocturnin* (*Noct*) is a prepachytene piRNA cluster (Li et al., 2013) that produces abundant piRNAs in BL6 and NOD strains but not in C3H and 129 (**Fig 1C, D and Supplementary Fig 2**). The gene *Zbtb37* produces significantly more piRNAs in strains NOD, C3H and 129 than in BL6 **(Fig 1C and Supplementary Fig 2).** The gene *Mrs2* produces piRNAs in three mouse strains but nearly none in NOD **(Fig 1C and Supplementary Fig 2).** Last, the two intergenic clusters on chromosome 14 produce piRNAs in all strains but significantly more piRNAs in NOD (**Fig 1C and Supplementary Fig 2**). The reference set of piRNA clusters for mouse corresponds to strain BL6 (Li et al., 2013). Thus, select piRNA clusters produce different steady state levels of piRNAs in mice of different strains.

We reasoned that there are likely additional piRNA clusters with significant expression differences between strains that are so far missed because they are not expressed in the BL6 strain. To address this bias for the reference mouse strain, we used the testis small RNA data from all four strains (BL6, NOD, C3H and 129) as well as eight samples from isolated spermatogonia from the reference mouse strain (BL6) and the wild-derived inbred mouse strain CAST/EiJ (referred to as CAST) to predict clusters *de novo* (see **Methods**). We then compared the expression of these predicted clusters between mouse strains (**Fig 2**). As expected, predicted piRNA clusters have more similar expression between samples of the same strain (**Fig 2A**). Also, piRNA cluster expression of samples from testis (enriched for pachytene piRNAs) and spermatogonia (enriched for prepachytene piRNAs) clustering separately (**Fig 2A**). Among the 845 predicted piRNA clusters, we found 93 that are differentially expressed in testis samples of the four strains. Thirty-five of these clusters are differentially expressed in testes of one of the four strains (i.e., significant in three pairwise comparisons) (**Fig 2B**). The samples from BL6 and CAST spermatogonia had the highest number of differentially expressed predicted clusters, likely because CAST is genetically more different from the rest of the classical inbred strains (**Fig 2C**). In total, we found 172 clusters differentially expressed in at least one of the pairwise strain comparisons and 59 clusters differentially expressed in at least three of the seven pairwise comparisons (six pairwise comparisons between testis samples from four strains and one pairwise comparison between spermatogonia samples from two strains) (**Supplementary Tables S5**, **S6**). Analysis of the data revealed predicted strain-specific uni-directional (e.g., **Fig 2D**) and bi-directional (e.g., **Fig 2E**) piRNA clusters expressed in testes, as well as strain-specific clusters expressed only in spermatogonia (e.g., **Fig 2F-G**). Thus, genomic loci exist within the mouse genome that produce significantly different amount of piRNAs in the germline of different strains of mice.

**Figure 2.**
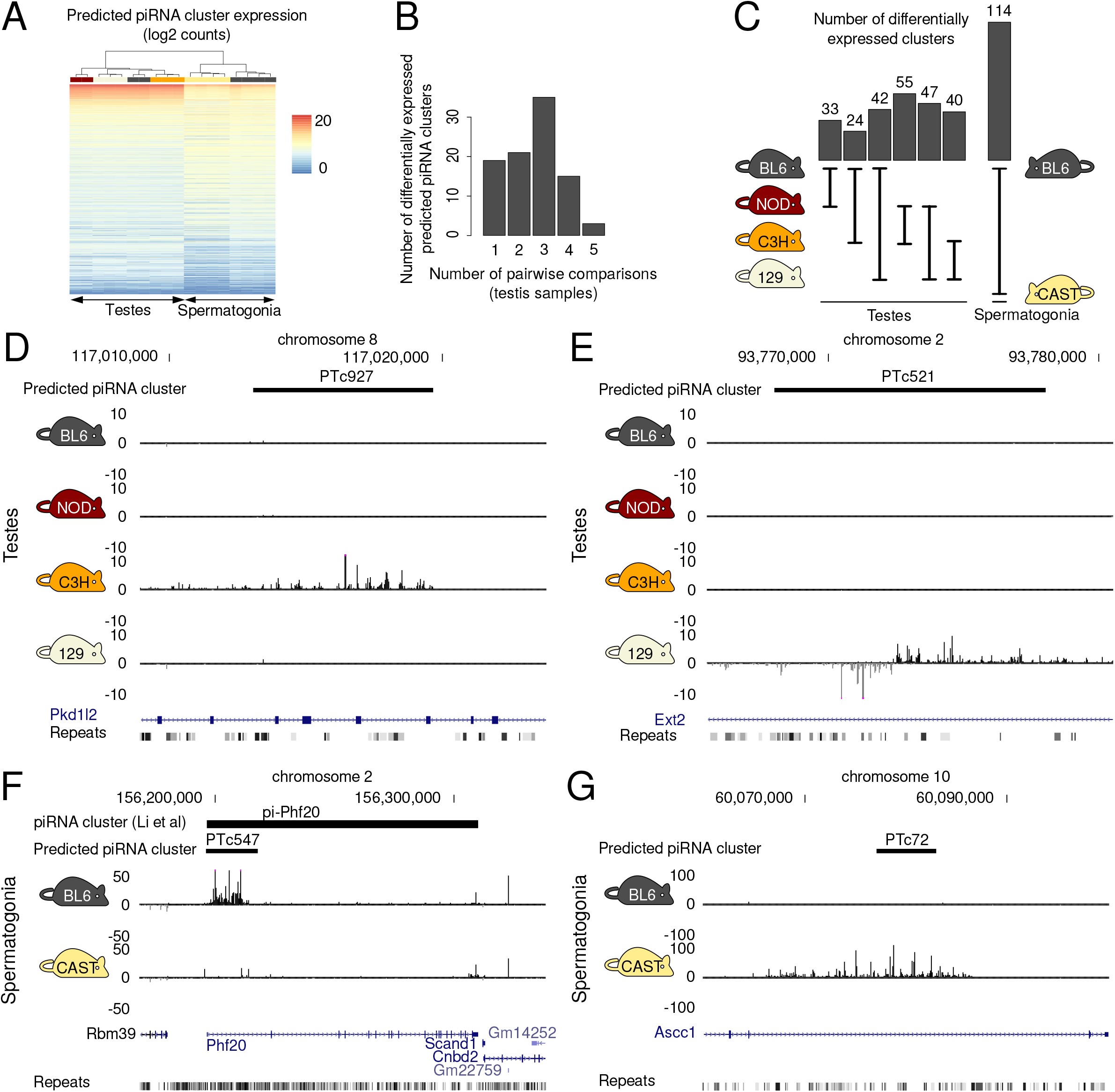
Many predicted piRNA-producing loci show significant differences in expression between mouse strains. (**A**) Heatmap showing clustering of expression of predicted piRNA-producing loci from whole testes and isolated spermatogonia from inbred mouse strains. (**B**) Frequency distribution of the number of differentially expressed predicted piRNA clusters in testis samples from four mouse strains. (**C**) Number of differentially expressed predicted piRNA-producing loci per pairwise strain comparison. (**D**) Uni-directional predicted piRNA cluster PTc927 is highly expressed only in testes of C3H mice. (**E**) Bi-directional predicted piRNA cluster PTc521 is highly expressed only in testes of 129 mice. (**F**) Predicted piRNA cluster PTc547 (cluster pi-Phf20 from Li et al 2013) is expressed in BL6 and not in CAST spermatogonia. (**G**) Predicted piRNA cluster PTc72 is expressed in CAST and not in BL6 spermatogonia. The positions of genes, repeats and multi-strain alignments from the UCSC Genome Browser are also shown.

### Association between an intronic IAP insertion and piRNA production from the mouse protein-coding gene *Nocturnin*

A locus with a notable difference in piRNAs between the four strains was the one overlapping the protein-coding gene *Nocturnin (Noct)*. *Noct* is one of the 114 previously annotated protein-coding genes that produce piRNAs in the mouse genome (Li et al., 2013). Considering small RNAs mapping uniquely to this locus, *Noct* produces a substantial number of piRNAs in testis samples of only two of the four strains (**Fig 3A**). We wondered what may be causing the apparent switch in production of piRNAs from this locus. Because transposons are tightly linked to the function and biogenesis of piRNAs, we turned our attention to known transposon insertions and deletions in the genomes of these strains (Nellåker et al., 2012). In the reference mouse strain, the first intron of *Noct* contains a 5.3kb ERV insertion of the IAP IΔ1 subtype and a small fragment of a LINE1 transposon that is variable between the five inbred strains (**Fig 3A**). The *Noct* IAP is a recent transposon insertion found in a subset of the laboratory inbred mouse strains(Dupressoir et al., 1999). Laboratory mouse strains have a common origin that dates back to the 1920s, making this insertion potentially less than a century old. Interestingly, mice of the two inbred strains that produce piRNAs from this locus (BL6 and NOD) carry the IAP insertion, while mice of the three strains that produce significantly fewer piRNAs (C3H, 129 and CAST) do not carry the insertion (**Fig 3A**). In contrast, production of piRNAs from the locus does not correlate with the presence of the variable LINE1 fragment, since CAST produces very few piRNAs but does not have the LINE1 fragment deletion (**Fig 3A**). The perfect correlation between the IAP insertion and pi-Noct abundance raises the possibility of a mechanistic link between IAP insertions and piRNA production.

**Figure 3.**
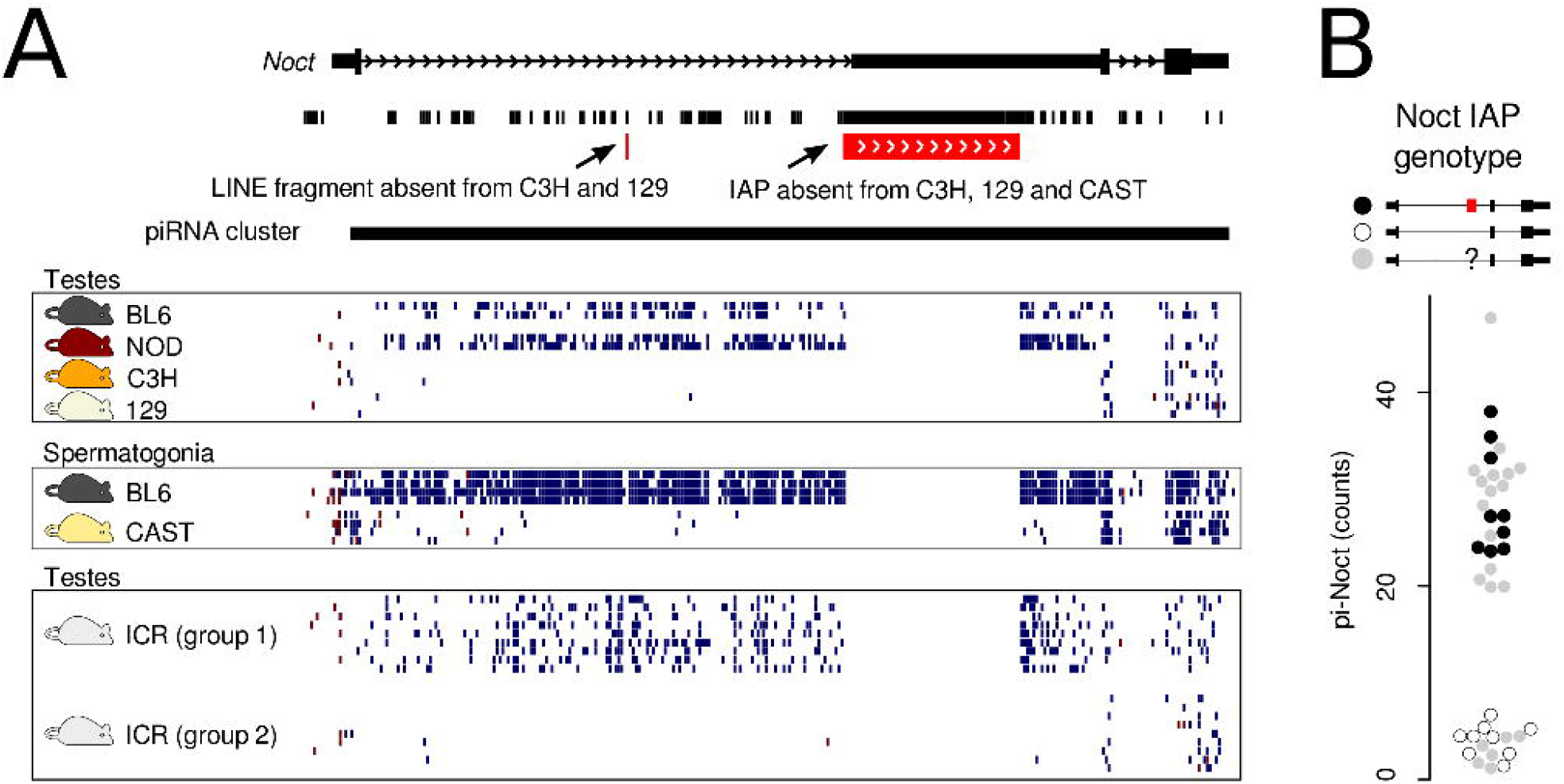
pi-Noct contains a polymorphic IAP that correlates perfectly with piRNA expression. (**A**) Mouse strains BL6, NOD produce many piRNAs mapping in the sense strand of the gene *Noct*, including its intron. *S*trains C3H, 129 and CAST produce negligible levels of small RNAs from the same genomic region. Nine samples from the outbred mouse strain ICR contain high levels of pi-Noct small RNAs (group 1), while another nine samples contain low levels (group 2). The eighteen ICR mice were genotyped, as shown in panel B. Only uniquely-mapping small RNAs are shown in this figure. Polymorphic transposable elements from (Nellåker et al., 2012) are indicated. (**B**) The data points show the normalised counts of small RNAs mapping to the pi-Noct locus in samples from 39 ICR mice. The data points fall into two groups according to their pi-Noct counts. Nine samples from each group were genotyped by PCR to test whether they contained the IAP. All nine samples with low pi-Noct expression were from mice without the IAP (shown as empty circles), while all nine samples with high pi-Noct expression were from mice with at least one copy of the IAP (shown as filled black circles). The rest of the samples are shown as filled grey circles.

Because inbred strains differ by many additional variants that are inherited together with the *Noct* IAP insertion and confound the association with piRNA abundance at this locus, we decided to analyse the expression of this piRNA cluster in mice from an outbred strain. We sequenced small RNAs from 39 young adult mice of the genetically outbred strain ICR (for further details on this dataset see **Methods** and **Supplementary Table 1**). In agreement with the results from the inbred strains, we found pi-Noct among the clusters with the highest variation in piRNA abundance (**Fig 3B, Supplementary Table S7**). To test the link between the variation in piRNA production from this locus and the presence of the IAP in *Noct*, we genotyped eighteen of these mice and confirmed the perfect association between piRNA production and the IAP insertion (**Fig 3B, Supplementary Figure 3** and **Methods**).

In summary, we found that genetic variation is linked to piRNA cluster expression in mouse. One of the loci with high piRNA abundance variation between animals overlaps the protein-coding gene *Noct*. The abundance of piRNAs produced from this locus in different animals perfectly agrees with the presence of the IAP in the first intron of the gene. These results suggest that the recent insertion of an IAP at this locus is mechanistically linked to piRNA production.

### General association between piRNA expression and transposable element variants

How pervasive is the association between new transposon insertions or deletions and variation in piRNA production? Although, transposons are depleted from genes (Nellåker et al., 2012) as well as from piRNA clusters (Aravin et al., 2006; Girard et al., 2006; Lau et al., 2006), some transposable element variants do overlap the predicted piRNA clusters. We used these annotated transposable element variants to test the association with piRNA cluster expression variation between mouse strains. Indeed, we found that clusters with significant differences in piRNA abundance between mouse strains are more common among clusters with transposable element variants than among the rest of the clusters (**Fig 4A**). Of three major types of transposons (LINEs, SINEs and ERVs), we found that the transposons with significant overrepresentation among predicted piRNA clusters with significant differences in piRNA expression between the strains were almost exclusively polymorphic ERVs, especially IAPs (**Fig 4A**). Clusters overlapping polymorphic IAPs are few, yet they include some of the known piRNA clusters with the biggest differences in piRNA abundance between any pair of the four mouse strains of this study (such as pi-Noct, 14-qA3-284, 14-qC1-1261 shown in **Fig 1**).

**Figure 4.**
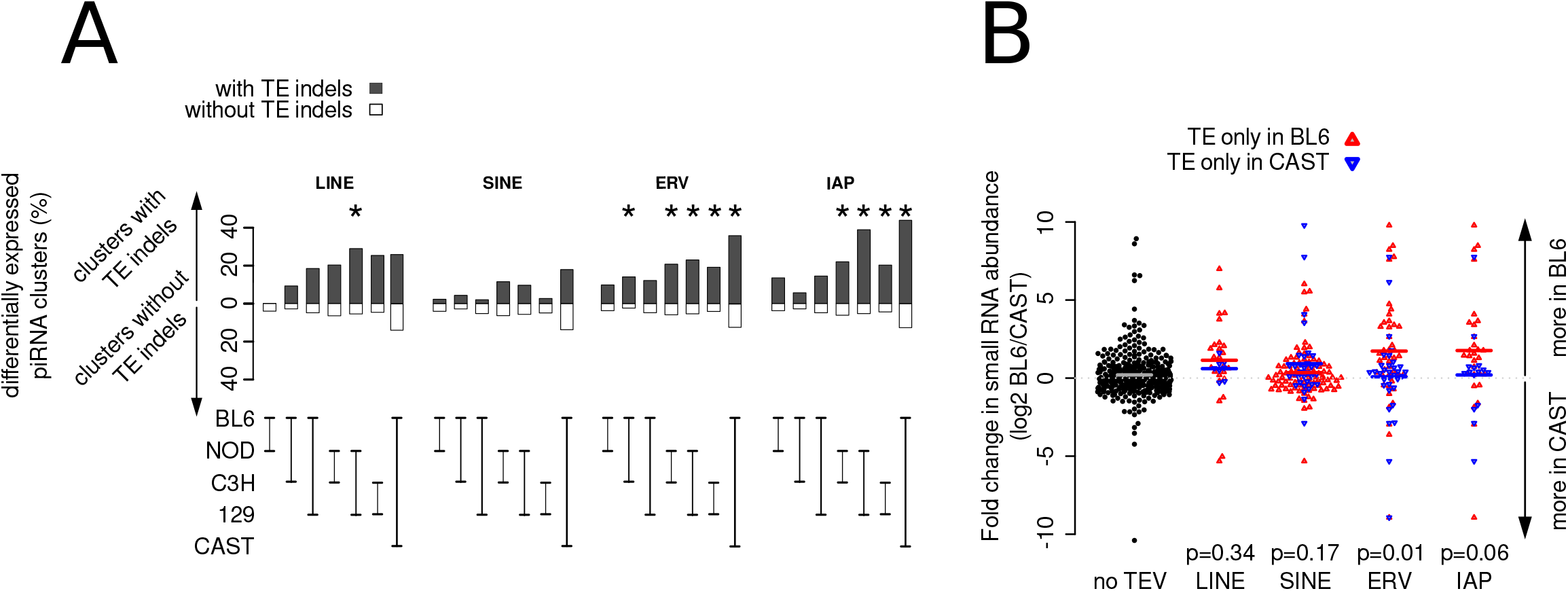
Polymorphic ERV insertions are significantly associated with highly variable piRNA production between mouse strains. (**A**) Percentages of predicted piRNA clusters with polymorphic transposable elements that are significantly differentially expressed in five strains are shown in black bars. Percentages of predicted piRNA clusters without polymorphic transposable elements are shown in white bars. Significant differences in these percentages, suggesting an association between differential expression and polymorphic transposable elements, were calculated using fisher’s exact test and are indicated with asterisks (p<0.05). (**B**) Predicted piRNA clusters with polymorphic ERVs have higher small RNA counts in the strain with the insertion. Polymorphic transposable elements were retrieved from (Nellåker et al., 2012). Data points showing fold-change in expression of clusters with insertions only in BL6 are indicated as upward facing, red triangles. Data points showing fold-change in expression of clusters with insertions only in CAST are indicated as downward facing, blue triangles. Data points showing fold-change in expression of clusters without polymorphic transposable elements (no TEV) are shown as filled, black circles. Coloured lines indicate group means. For each transposon class, we tested that the distribution of changes in piRNA expression between BL6 and CAST is the same for clusters with transposable elements that are only found in BL6 (red points) as for clusters with transposable elements only in CAST (blue data points), using the Wilcoxon rank-sum test in R (p-values shown).

We tested whether transposable element insertions are associated with an increase in piRNA abundance, as seen for pi-Noct. We focused on the comparison between BL6 and CAST because this strain pair has both the highest number of different ERV variants and the highest number of differentially expressed predicted piRNA clusters (**Fig 2C**). We split the 74 predicted clusters with variable ERVs between these strains into those with the transposable element only in BL6 (**Fig 4B**, data points shown as red triangles pointing up) and those with the transposable element only in CAST (**Fig 4B**, data points shown as blue triangles pointing down). We found that the abundance of piRNAs from clusters with ERVs in BL6 was higher in BL6 than in CAST and vice versa (Wilcoxon-rank-sum test p-value = 0.01). The same trend can be seen for clusters with IAP variants but without passing the significance threshold (Wilcoxon-rank-sum test p-value = 0.06), likely due to the lower total count of clusters with IAP variants (44 clusters). We did not find a significant association between clusters with insertions or deletions of LINEs or SINEs and the direction of piRNA abundance change between the two strains (**Fig 4B**). These observations are not due to the expression of the repetitive element itself or an artefact of ambiguous mapping of small RNA data to the mouse genome, since the changes in small RNA abundance that we report here are based on expression values calculated only from uniquely mapping small RNA reads that align outside annotated repeats. Taken together, the data suggests that ERV insertions can cause an increase of piRNA production or expression from diverse genomic loci in mouse. ERV insertions could trigger the emergence of novel piRNA clusters during evolution.

### The IAP insertion in *Noct* is associated with post-transcriptional processing of germline-expressed transcripts into piRNAs

IAPs can affect gene expression in multiple ways, one of which is by regulating transcription. Thus, we asked whether piRNA production is explained by IAP-induced ectopic transcriptional activation of the gene during spermatogenesis. *Noct* is a gene known to be expressed in many mouse organs, including testes (Dupressoir et al., 1999). Still, the relative expression of the different *Noct* alleles during spermatogenesis had not been studied. To address this, we analysed available steady state gene expression data from various stages of spermatogenesis from 129/DBA hybrid mice (Gan et al., 2013) carrying one *Noct* allele with the IAP insertion (inherited from the DBA parent) and the second allele without (inherited from the 129 parent). Using single nucleotide polymorphisms specific to each of the parental strains, we quantified the expression of the two different alleles in 129/DBA mouse male germ cells and tested whether the *Noct* allele carrying the IAP insertion is more abundantly expressed than the other allele. We found that throughout spermatogenesis *Noct* is very highly expressed (**Fig 5A**, lower panel) with no evidence of the *Noct* allele with the IAP being more highly expressed than the *Noct* allele without the IAP (**Fig 5A**, upper panel). Similarly, we analysed the chromatin state of *Noct* using available H3K4me3 ChIP-seq data from spermatocytes of mouse BL6/CAST hybrid mice (Baker et al., 2015) carrying one *Noct* allele with the IAP insertion (BL6) and another without (CAST). Again, we found that the only active promoter region along the gene is that of the actual *Noct* promoter, with no evidence of additional H3K4me3-marked regions surrounding the IAP, and with the two alleles showing no differences in terms of this active chromatin mark (**Fig 5B**). These results argue that the IAP inserted into an existing germline-expressed gene during very recent murine evolution and that this insertion did not discernibly alter transcription.

**Figure 5.**
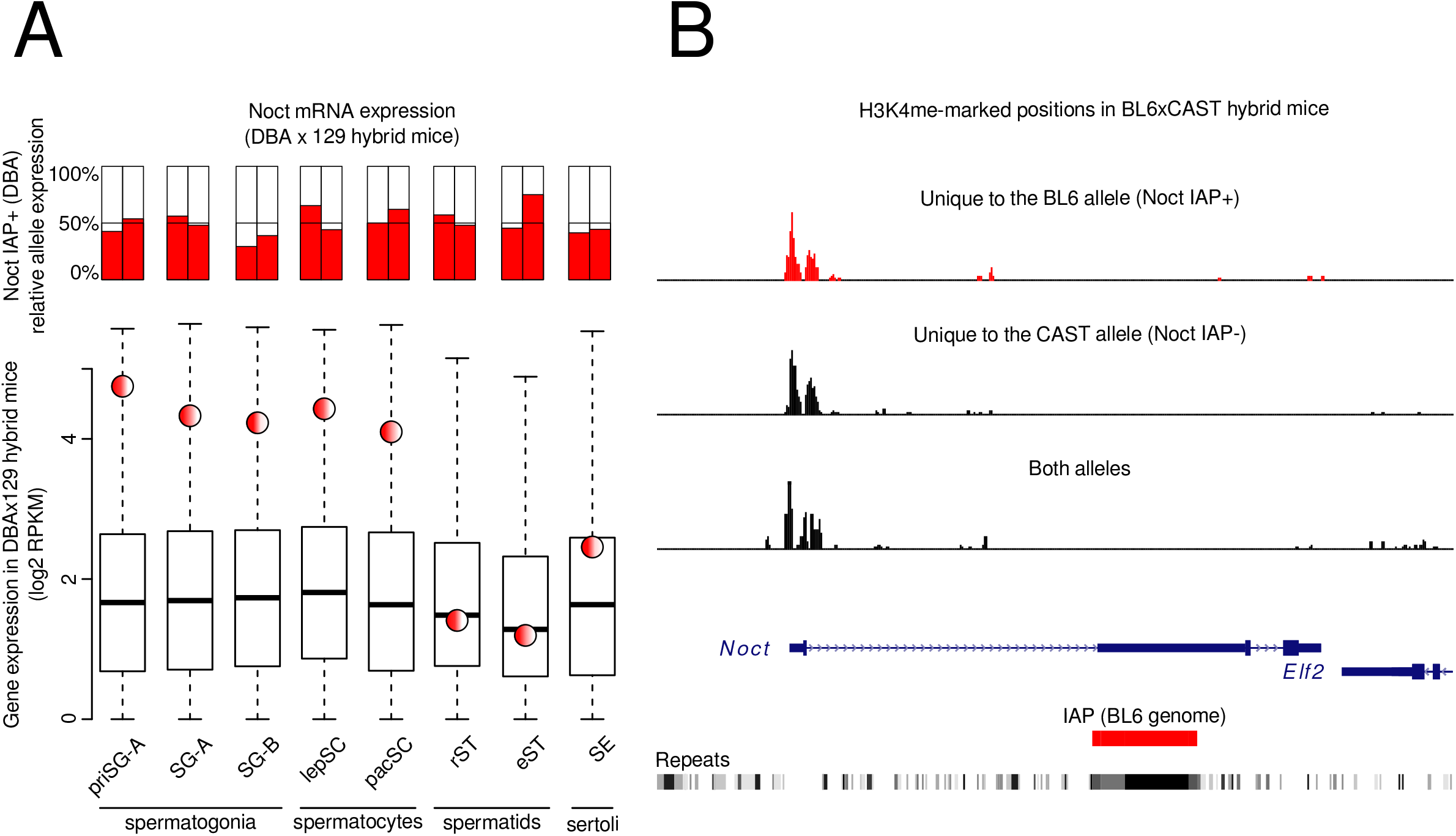
No evidence of differential mRNA expression due to the presence of the IAP insertion in *Noct*. (**A)** The bar plots show the relative expression of two different *Noct* alleles in 129xDBA F1 hybrid mice carrying one allele with the IAP (inherited from the DBA father) and one without (inherited from the 129 mother). The side-by-side pairs of barplots correspond to biological replicates. The y-axis shows the percentage of strain-specific RNA-seq reads that map to the DBA *Noct* allele (IAP+). The box plots show the distribution of expression of all genes during spermatogenesis in 129xDBA F1 hybrid mice. The red and white circle shows the expression of *Noct* in each cell type. **(B)** Allele-specific analysis of H3K4me3 ChIP-Seq from BL6xCAST F1 hybrid mice shows that the active H3K4me3 chromatin mark is found at similar levels on both *Noct* alleles. Reads mapping unambiguously to each of the two alleles using strain-specific single nucleotide polymorphisms (SNPs) are shown on the top two tracks. Uniquely mapping reads that do not overlap strain-specific SNPs are shown in the bottom track.

An alternative mechanism that could explain the observed data is that the IAP carries a signal involved in post-transcriptional regulation that induces piRNA production from transcripts. This signal is not just sequence complementarity between an antisense piRNA matching the IAP inside the *Noct* primary transcript, because this would trigger piRNA production only downstream of the IAP. As shown in Fig1C and 3A, at this locus, piRNAs are also produced upstream of the IAP, most likely from the primary unspliced transcript transcribed from the first *Noct* promoter. Thus, in this case, it looks like the IAP is causing the unspliced transcript to be exported from the nucleus and to be recognised as a piRNA precursor. We tested whether the association between polymorphic IAP insertions and piRNA production depends on the orientation of the IAP. Comparing small RNA production from predicted piRNA clusters from BL6 and CAST spermatogonia, we found no difference in piRNA levels at loci with strain-specific ERVs antisense to the piRNA cluster (**Fig 6**). Importantly, however, strong and significant associations were observed specifically where ERV insertions are in the piRNA producing strand (**Fig 6**). Thus, we conclude that ERV insertions can trigger and/or enhance piRNA production from existing transcribed genomic loci, likely through a post-transcriptional mechanism and that this mechanism appears to require the ERV to be oriented in sense to the host transcript.

**Figure 6.**
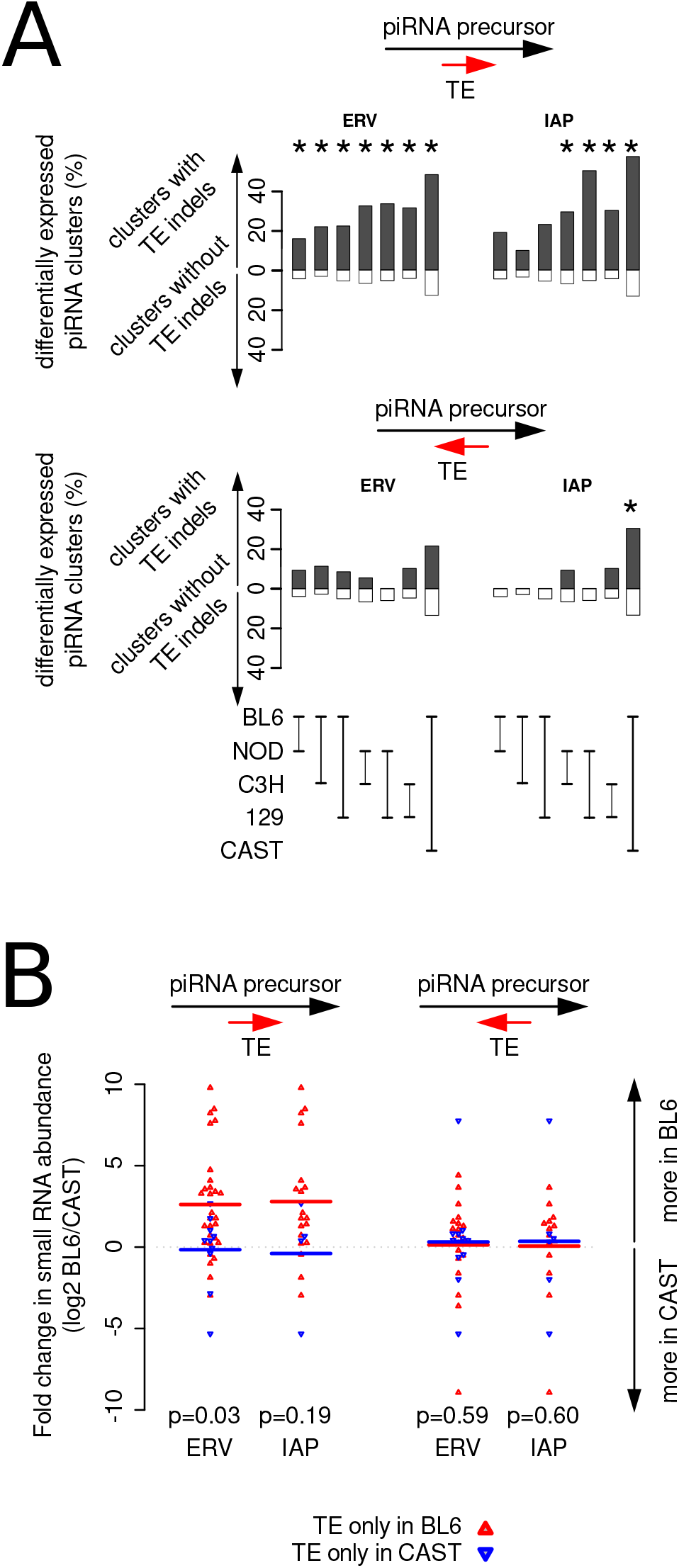
ERV insertions specifically on the sense strand of the precursor transcript are associated with higher piRNA production between mouse strains. (**A**) Predicted piRNA clusters with ERV insertions in the sense strand (upper panels) and in the antisense strand (lower panels) were analysed separately. Percentages of predicted piRNA clusters with polymorphic transposable elements that are significantly differentially expressed in five strains are shown in black bars. Percentages of predicted piRNA clusters without polymorphic transposable elements are shown in white bars. Significant differences in these percentages, suggesting an association between differential expression and polymorphic transposable elements, were calculated using fisher’s exact test and are indicated with asterisks (p<0.05). (**B**) Predicted piRNA clusters with polymorphic ERVs have higher small RNA counts in the strain with the insertion, only when the insertion is in the same strand as the predicted cluster. Polymorphic transposable elements were retrieved from (Nellåker et al., 2012). Data points showing fold-change in expression of clusters with insertions only in BL6 are indicated as upward facing, red triangles. Data points showing fold-change in expression of clusters with insertions only in CAST are indicated as downward facing, blue triangles.

## Discussion

We uncovered significant variation in piRNA production from a subset of piRNA-producing loci in genetically diverse mouse strains. This is the first comparison of piRNA production in different animals of any mammalian species. Our results are in agreement with what was previously observed in different strains of flies (Song et al., 2014). In addition to quantitative differences in piRNA production, both mice and flies have strain-specific sources of piRNAs. Therefore, the rapid emergence of multiple new piRNA-producing loci within a species is a core property of the piRNA system and likely to be found in all animals expressing this pathway. The high within-species diversity also agrees with the high between-species divergence of piRNA-producing loci in animals (Assis & Kondrashov, 2009; Chirn et al., 2015).

One of the primary mechanisms of novel piRNA production in a species appears to be the insertion of transposable elements to new positions in the genome. In mice, we found a significant association between piRNA production and novel transposable element insertions, in particular IAPs. In flies, strain-specific piRNA clusters were found at positions of novel insertions of LTR and LINE elements (Mohn et al., 2014; Shpiz et al., 2014; Song et al., 2014). Our analysis revealed that transposable element insertions or deletions are often - but not always - found at clusters that show major inter-strain differences in piRNA production. There are several possible explanations for this. It is possible that the annotation of transposable element variants is incomplete, and all strain-specific piRNA-producing loci are due to transposable elements insertions and deletions. Additionally, some of the differences in piRNA abundance could be due to differences in the expression level or the processing of the precursor. For example, genetic variation leading to the gain of a new binding site for A-Myb, the major transcription factor for pachytene piRNA expression, could explain the birth of some of the strain-specific piRNA clusters. Differences in the steady state level of the precursor was not the case for one of the highly variable piRNA clusters that we studied in detail (pi-Noct), but it may be the case for others.

We found that the orientation of the ERV insertion appears coupled to inter-strain piRNA cluster expression differences. In particular, we found that ERV insertions only had an effect when they were in sense to the piRNA precursor. The IAP insertion in the first intron of the gene *Noct* fits this model. In the case of pi-Noct we found that the IAP insertion does not modify the expression level of the precursor. It is also clear that piRNA production from *Noct* by antisense piRNAs targeting the IAP is not the mechanism that leads to piRNA production from this locus. This is because most piRNAs are produced from the intron, upstream of the position of the IAP insertion. The fact that piRNAs are produced from the intron, also highlights the fact that *Noct* transcripts producing piRNAs are either unspliced or aberrantly spliced, that they evade surveillance mechanisms in the nucleus and that they are exported to the cytoplasm, where piRNA biogenesis takes place. The fact that IAP-containing, unspliced *Noct* transcripts produce piRNAs suggests that these transcripts are processed into piRNAs because they are recognised as retroviruses, like KoRV-A in Koala and the AKV Murine Leukemia Virus in the AKR mouse strain (Yu et al., 2019). However, unlike KoRV-A and AKV elements, the IAP is embedded within the intron of a gene and transcribed with it. It reveals how an ERV insertion in the intron of a protein-coding gene can signal a much greater transcript for piRNA biogenesis. We can only speculate about the mechanism of piRNA production from the IAP-containing allele of *Noct*. It could be the absence of splicing, as previously proposed by Yu et al, that signals that this transcript should be sliced into piRNAs. It could also be the presence of a strand-specific signal within the IAP interacting with a nuclear exporter and piRNA biogenesis factor. Functional experiments are necessary to further dissect the mechanism by which the *Noct* IAP insertion leads to piRNA production from this locus.

We currently do not know whether the differences in piRNA content between animals of the same strain have biological or physiological consequences, conferring higher or lower fitness. We speculate that the burst of novel piRNAs triggered by a transposon insertion event has the potential to generate new regulatory effects in *cis* and in *trans*. As with other genetic variants, the emergence of new piRNAs can be beneficial (recognition of invading parasitic elements) or deleterious (silencing of essential protein-coding genes) for the organism. What is unique to polymorphic piRNA producing loci is the magnitude of new material for natural selection to act upon. It is perhaps because of the many possibilities for positive or negative effects on fitness by each individual piRNA, among the many produced from a single locus, that piRNA-producing loci are gained and lost so fast during evolution.

In conclusion, by sequencing and analysing small RNAs from the male germlines of different mouse strains we identified polymorphic and variably expressed piRNA-producing loci in a mammalian species. Insertions and/or deletions of active ERVs at germline-expressed genomic loci are two genetic mechanisms that spark piRNA expression variation and diversity, but there are certainly more to be discovered. Although small RNA data from inbred mouse strains were essential for the documentation of within-species differences in piRNAs determined by genetics, they could not be used to identify genetic variants associated with piRNA cluster expression in a global, systematic and unbiased way, because classical inbred strains vary at millions of positions along their genomes (Keane et al., 2011). Nevertheless, the data we generated chart the degree of piRNA cluster expression variation to be expected between genetically different animals of this rodent species. Some mouse strains have very young copies of IAPs consistent with activity in recent years (Nellåker et al., 2012). It remains to be seen whether ERV insertions and deletions are also a significant source of piRNA expression variation and diversification in other mammalian species, such as human.

## Materials and Methods

### Mouse tissue isolation and RNA extraction

Testes used in this study were obtained from mice from various sources, all following institutional regulations for animal care and use. Specifically, ICR (ICR-CD1, Envigo) and 129S1/SvlmJ (local established colony from previously purchased animals from The Jackson Laboratory) mice were maintained and used according to the guidelines of the Universitat de Barcelona Animal Care and Use Committee, C3HeB/FeJ and NOD mice were maintained and used according to the guidelines of the animal facility of the Institute Germans Trias I Pujol research institute (IGTP). All testis used in this work were from young adult mice. Testicles were rapidly dissected, snap-frozen in liquid nitrogen and stored at −80° C. For the sequencing of small RNA from classical inbred mouse strains, total RNA was extracted from previously frozen testes using TRIzol Reagent (Life Technologies; Thermo Fisher Scientific) linked to PureLink RNA Mini Kit (Invitrogen: Thermo Fisher Scientific) following the manufacturer’s protocol “TRIzol Plus Total Transcriptome Isolation”.

For the isolation of spermatogonial RNA, C57BL/6J and Cast/EiJ mice were obtained from The Jackson Laboratory and kept in the SPF animal facility of Max Planck Institute of Immunobiology and Epigenetics until sacrifice. In order to isolate spermatogonia from mice, testes were dissected and digested according to the protocol by Liao et al ((H.-F. Liao et al., 2016), with minor modifications. Briefly, we euthanized 6 weeks old mice with CO_2_ and quickly dissected testes, removed the tunica albuginea and loosened the seminiferous tubules. We then digested these tissues with 1 mg/ml collagenase IV (Worthington, LS004189) in DMEM (Gibco, 31966-024) supplemented with 10% FBS, 100 U/ml penicillin-streptomycin (Gibco, 15140-122), 250 ng/ml fungizone (Gibco, 15290-018) and 50 μg/ml gentamycin (Serva, 4799.01) in a petri dish at 37°C over a Thermoblock, shaking at 600 rpm for 30 minutes. The reaction continued for another 10 minutes after the addition of 0.25% tripsin EDTA (Sigma, T4849) at 37°C and 600 rpm. We homogenized the digested tissues by pipetting up and we washed the solution with a double amount of PBS (Gibco, 14190-094) supplemented with 10% FBS. Pieces of remaining, undigested tissues were filtered with a 40 μm strainer (BD Falcon, 352340). The filtered solution was then centrifuged at 300 g for 10 minutes at 4°C. We removed the supernatant and then resuspended the pellet in 200 μl of FACS buffer (PBS supplemented with 5% BSA and 5 mM EDTA) supplemented with 1U/μl SUPERase.in (Invitrogen, AM2696). Spermatogonia were sorted according to (Kanatsu-Shinohara et al. 2011) for the expression of CD9 (eBioscience, 17-0091-82, 1μg) and Epcam (eBioscience, 0.125 μg). Sorted cells were centrifuged at 300g for 10 minutes at 4°C and resuspended in 1 ml of TRIzol (Invitrogen, 15596018). Spermatogonial RNA was purified according to the standard TRIzol protocol and contaminant genomic DNA was digested using the DNA-free kit (Invitrogen, AM1906).

### Small RNA-seq library preparation and sequencing

All small RNA sequencing libraries were prepared by the Genomics and Bioinformatics Facility of the IGTP. Libraries were prepared with TruSeq small RNA from illumina with extended range of size selection. Pippin prep was used for automated pooled library size selection. Libraries were indexed using Illumina barcodes and sequenced using a HiSeq2500 (Illumina) as single 50nt reads. Small RNA libraries corresponding to samples from inbred strains were sequenced as a single pool on two lanes and the resulting data (all showing very high correlation between lanes) were merged for analysis.

### Small RNA-seq data analysis

We removed the adapter (TGGAATTCTCGGGTGCCAAGGAACTCCAGTCAC) from the small RNA reads using cutadapt v2.10 (Martin 2011), requiring 9nt of match with the adapter. We discarded reads shorter than 19nt, longer than 36nt and any reads not matching the adapter. We filtered reads based on quality using the FASTX Toolkit v0.0.14 (http://hannonlab.cshl.edu/fastx_toolkit/) allowing minimum quality score 30 over at least 90% of nucleotides. We then mapped the trimmed and filtered reads to the reference mouse genome (primary assembly GRCm38/mm10) using bowtie v1.2.3(Langmead et al., 2009) with the options -M 1 --best --strata −v 1 to get the best alignment with up to 1 mismatch, reporting only one match for multimapping reads.

### Prediction of piRNA producing loci

We used the proTRAC pipeline v2.4.4 (Rosenkranz & Zischler, 2012) to predict clusters for each of the 18 samples from inbred mice with default options. To get a set of predicted piRNA clusters for each strain and sample type, we took the intersection of the clusters predicted using the samples of each strain. This resulted in four sets of predicted clusters for testis samples (BL6, NOD, C3H, 129) and two sets of predicted clusters for spermatogonia samples (BL6 and CAST). To get one list of predicted clusters for mouse, we merged the coordinates of the six sets. From this set, we removed clusters that matched repeats (RepeatMasker annotation) and polymorphic transposable elements by more than 80% of their length resulting in a set of 981 mouse predicted clusters. This set included regions with overlapping clusters predicted to be bidirectional in some samples and unidirectional in others. To avoid double counting of regions during differential cluster expression, we removed those predicted bidirectional promoters that overlapped unidirectional promoters, reducing the total set to 865 predicted piRNA clusters. Finally, we removed clusters with total read count of less than 10 in our entire dataset. The final set consists of 845 predicted piRNA clusters. The coordinates of these regions and the results of differential expression in testis of four inbred strains and in spermatogonia of two inbred strains are provided in Supplementary Tables 5 and 6, respectively.

### Differential Expression Analysis of piRNA producing loci and test of association with variable transposable elements

To quantify the expression of piRNA clusters (known or predicted), we annotated reads mapping to the clusters using featureCounts v2.0.1 (Y. Liao et al., 2014) with the options -Q 1 −s 0 -minOverlap 18 to count reads with minimum quality score of 1, mapping within the region of the cluster with a minimum overlap of 18nt. To reduce possible artefacts due to differences in repeat content or repeat expression between strains, for differential expression analysis we only counted reads mapping to unique locations in the reference mouse genome and only reads not overlapping repeats from the RepeatMasker annotation of the reference mouse genome. The same analyses without removing reads mapping to annotated repeats were qualitatively similar (data not shown). We removed from differential expression analysis clusters with fewer than ten reads in all samples. Predicted clusters overlapping (for example in cases where a unidirectional and a bidirectional cluster overlapped) were removed from statistical tests. For differential piRNA cluster expression analysis we used DESeq2 v1.34.0(Love et al., 2014) with absolute log_2_ fold change threshold great than 1 and false discovery rate threshold of 0.05.

We retrieved the annotation of variable transposable elements from Additional file 13 from Nellaker et al(Nellåker et al., 2012). We grouped the retrieved variable transposable elements into SINEs (as annotated in the file), LINEs (annotated as LINE and LINE fragments), ERVs (the rest of the elements in the retrieved file, which are different families of ERVs) and IAPs (annotated as IAP-I). For association with predicted piRNA clusters, we considered that predicted clusters overlapped variable transposable elements if they were within 5kb from one and tested the association using Fisher’s exact test in R with significance threshold of 0.05. For tests of association using strand information, we only considered clusters and repeats with annotated strand as + or - (bidirectional piRNA clusters were excluded from this analysis). The significance of the differences in the distribution of fold change expression of predicted clusters in different strains (Fig 6B) was using the two-sample Wilcoxon rank sum test in R.

### RNA-seq and ChIP-Seq allele-specific data analysis

To test for differences in expression of *Noct* IAP+ and *Noct* IAP- alleles, we retrieved RNA-seq data from GSE35005 (Gan et al., 2013) from DBA/2NCrlVr x 129S2/SVPasCrlVr F1 hybrid mice. According to the annotated variable transposable elements of eighteen genotyped mouse strains(Nellåker et al., 2012), three 129 strains (129S1/SvImJ, 129P2/OlaHsd and 129S5/SvEvBrd) carry the IAP insertion in *Noct.* We thus expect that 129S2/SVPasCrlVr also carries it. As noted in Nellaker et al, mouse substrains are nearly identical to each other in comparison to other strains. Similarly, following the same line of thought we expect that the DBA/2NCrlVr strain carries the same *Noct* allele as the genotyped strain DBA/2J. To test for differences in the H3K4me3 chromatin mark on a *Noct* IAP+ and a *Noct* IAP- allele, we retrieved ChIP-seq data from GSE60906 (Baker et al PLoS Genet 2015) from C57BL/6J x CAST/EiJ F1 hybrid mice. According to the genotyped mouse strains, C57BL/6J carries the *Noct* IAP+ allele and CAST/EiJ carries the *Noct* IAP- allele.

To retrieve reads mapping to the two different alleles in the samples of the hybrid mice, we used SNPSplit v0.3.3 (Krueger & Andrews, 2016). Briefly, we masked the reference mouse genome changing all the SNP positions to Ns. The list of SNPs between mouse strains and the reference mouse genome was retrieved from the Sanger Institute Mouse Genomes Project v5, dbSNP142. RNA-seq reads were mapped using HISAT2 v2.2.1(Kim et al., 2019) with options -no-softclip using known splice sites from the reference mouse genome. ChIP-seq reads were mapped using bowtie2 v2.2.5(Langmead et al., 2009) with default options. Reads overlapping SNPs were assigned to the corresponding strain using SNPSplit.

### Noct allele genotyping of ICR mice

For genotyping the *Noct* IAP (Fig 3B and Supplementary Figure 3), DNA extraction from mouse liver tissue was performed using the Maxwell 16 Tissue DNA Purification kit (Promega) following the manufacturer’s protocol. 20 μL PCR reactions were performed with 50 ng of genomic DNA using the Phusion High Fidelity DNA Polymerase (2 U/μL) (Life Technologies) following manufacturer’s indications. Specifically, 0.5 μL of 10 μM Forward primer (5’ TACTAATTCCAGACCTCTCTCC 3’) and Reverse primer (5’ GCACTGTAGAGTCGACTGGTGC 3’) were used together with 0.4 μL 10 mM dNTPs and 0.4 μL of Phusion Polymerase. PCR conditions were as follows: an activation step at 98°C for 3’; 30 x 3-step cycles of denaturing at 98°C for 10’’, annealing at 61.2°C for 20’’ and extension at 72°C for 4’ 15’’; followed by a final step at 72°C for 5’. Amplicons were run in 0.8% agarose gels stained with SYBR safe (Life Technologies). Gel pictures were taken with Molecular Imager^®^ Gel Doc™ XR+ imaging system (BioRad).

## Acknowledgements

We thank Pere-Joan Cardona from the Microbiology Service at Hospital Germans Trias i Pujol and Jorge Díaz, from the CMCiB for kindly providing tissues of the C3HeB/FeJ mouse strain, Marta Vives-Pi from the Immunology of Diabetes Unit at Germans Trias i Pujol Research Institute (IGTP) for tissues of the NOD mouse strain and Jordi Llorens from the Department of Physiological Sciences, University of Barcelona for tissues of the 129S1/SvlmJ strain. We thank Lauro Sumoy and the High Content Genomics and Bioinformatics Facility of the IGTP for small RNA library preparation, the IGTP and the IJC High Performance Computing Core Facilities for systems administration and the CRG Genomics Unit for small RNA sequencing. We thank Ben Lehner, Adelheid Lempdradl and Anita Ost for comments on an earlier version of the manuscript. EC was funded by an AGAUR FI PhD fellowship. This work was funded by Grant BFU2015-70581 and PID2019-111676GB-I00 from the Spanish Ministry of Economy and Competitiveness and the CERCA Programme/Generalitat de Catalunya.

## Author contributions

Conceived and designed the experiments: EC SF TV. Performed the experiments: EC PS CM JC IP SF TV. Analysed the data: EC PS CM SF TV. Contributed reagents/materials/analysis tools: EC PS CM IP JC. Wrote the paper with input from all authors: EC PS TV. Supervised: TV AP SF JC.

## Data and code availability

All sequencing data produced for this publication have been deposited in the NCBI Gene Expression Omnibus under accession number GSE215030. Custom scripts written to analyse the data are available from https://github.com/vavouri-lab/mouse-piRNA-variation.

## Supplementary Figures

**Supplementary Figure S1.** Abundance of piRNAs from prepachytene, pachytene and hybrid piRNA cluster in whole testis samples of the four inbred mouse strains. The classification of the three sets of clusters were retrieved from (Ding et al., 2018).

**Supplementary Figure S2.** Classical inbred mouse strains produce significantly different levels of piRNAs from pi-Noct (also known as pi-Ccrn4l) (A), pi-Zbtb37 (B), pi-Mrs2 (C), 14-qA3-284 (D) and 14-qC1-1261(E). Protein-coding genes and repeats at these loci are also shown.

**Supplementary Figure S3.** Genotyping of *Noct* IAP in mice. The top panel shows the position of the primers used for genotyping PCR. The IAP-containing *Noct* allele produces a PCR amplicon that is 8037bp long while the *Noct* allele without the IAP produces an amplicon that is 2720bp. Uniquely mapping small RNAs from a representative ICR mouse sample (with high pi-Noct expression) are shown in red. All small RNAs map in the sense strand. Samples from inbred mouse strains with the *Noct* IAP insertion (BL6 and NOD) and without the *Noct* IAP insertion (129 and C3H) were included as controls. The samples used for the genotyping were from the same mice as those used to generate small RNA sequencing data (labelled sample01-sample06 and sample16-sample27 in Supplementary Table S1)

## Supplementary Tables

**Supplementary Table S1.** Small RNA sequencing summary.

**Supplementary Table S2.** Raw small RNA counts in testis samples from inbred mouse strains for 214 previously described piRNA clusters.

**Supplementary Table S3.** Normalised small RNA counts in testis samples from inbred mouse strains for 214 previously described piRNA clusters.

**Supplementary Table S4.** Differential piRNA cluster expression in testis samples from four inbred strains for 214 previously described piRNA clusters.

**Supplementary Table S5.** Differential predicted piRNA cluster expression in testis samples from four inbred strains.

**Supplementary Table S6.** Differential predicted piRNA cluster expression in spermatogonia samples from BL6 and CAST strains.

**Supplementary Table S7.** Normalised small RNA counts in testis samples from ICR mice for 214 previously described piRNA cluster (see Table S1 for details on these samples).

**Supplementary Table S8.** Significance of differential expression of predicted piRNA clusters and overlap with transposable element variants between the five inbred mouse strains.

## Notes

### Competing Interest Statement

The authors have declared no competing interest.

## References

Aravin, A., Gaidatzis, D., Pfeffer, S., Lagos-Quintana, M., Landgraf, P., Iovino, N., Morris, P., Brownstein, M. J., Kuramochi-Miyagawa, S., Nakano, T., Chien, M., Russo, J. J., Ju, J., Sheridan, R., Sander, C., Zavolan, M., & Tuschl, T. (2006). A novel class of small RNAs bind to MILI protein in mouse testes. Nature, 442(7099), 203–207. https://doi.org/10.1038/nature04916

Assis, R., & Kondrashov, A. S. (2009). Rapid repetitive element-mediated expansion of piRNA clusters in mammalian evolution. Proceedings of the National Academy of Sciences of the United States of America, 106(17), 7079–7082. https://doi.org/10.1073/pnas.0900523106

Baker, C. L., Kajita, S., Walker, M., Saxl, R. L., Raghupathy, N., Choi, K., Petkov, P. M., & Paigen, K. (2015). PRDM9 drives evolutionary erosion of hotspots in Mus musculus through haplotype-specific initiation of meiotic recombination. PLoS Genetics, 11(1), e1004916. https://doi.org/10.1371/journal.pgen.1004916

Chirn, G., Rahman, R., Sytnikova, Y. A., Matts, J. A., Zeng, M., Gerlach, D., Yu, M., Berger, B., Naramura, M., Kile, B. T., & Lau, N. C. (2015). Conserved piRNA Expression from a Distinct Set of piRNA Cluster Loci in Eutherian Mammals. PLOS Genetics, 11(11). https://doi.org/10.1371/journal.pgen.1005652

Cosby, R. L., Chang, N.-C., & Feschotte, C. (2019). Host–transposon interactions: conflict, cooperation, and cooption. Genes & Development, 33(17–18), 1098–1116. https://doi.org/10.1101/gad.327312.119

Ding, D., Liu, J., Midic, U., Wu, Y., Dong, K., Melnick, A., Latham, K. E., & Chen, C. (2018). TDRD5 binds piRNA precursors and selectively enhances pachytene piRNA processing in mice. Nature Communications, 9(1), 127. https://doi.org/10.1038/s41467-017-02622-w

Dupressoir, A., Barbot, W., Loireau, M.-P., & Heidmann, T. (1999). Characterization of a Mammalian Gene Related to the Yeast CCR4 General Transcription Factor and Revealed by Transposon Insertion. Journal of Biological Chemistry, 274(43), 31068–31075. https://doi.org/10.1074/jbc.274.43.31068

ElMaghraby, M. F., Andersen, P. R., Pühringer, F., Hohmann, U., Meixner, K., Lendl, T., Tirian, L., & Brennecke, J. (2019). A Heterochromatin-Specific RNA Export Pathway Facilitates piRNA Production. Cell, 178(4), 964–979.e20. https://doi.org/10.1016/j.cell.2019.07.007

Gainetdinov, I., Colpan, C., Arif, A., Cecchini, K., & Zamore, P. D. (2018). A Single Mechanism of Biogenesis, Initiated and Directed by PIWI Proteins, Explains piRNA Production in Most Animals. Molecular Cell, 71(5). https://doi.org/10.1016/j.molcel.2018.08.007

Gan, H., Wen, L., Liao, S., Lin, X., Ma, T., Liu, J., Song, C., Wang, M., He, C., Han, C., & Tang, F. (2013). Dynamics of 5-hydroxymethylcytosine during mouse spermatogenesis. Nature Communications, 4(1), 1995. https://doi.org/10.1038/ncomms2995

Girard, A., Sachidanandam, R., Hannon, G. J., & Carmell, M. A. (2006). A germline-specific class of small RNAs binds mammalian Piwi proteins. Nature, 442(7099), 199–202. https://doi.org/10.1038/nature04917

Kaaij, L. J. T., Hoogstrate, S. W., Berezikov, E., & Ketting, R. F. (2013). piRNA dynamics in divergent zebrafish strains reveal long-lasting maternal influence on zygotic piRNA profiles. RNA, 19(3), 345–356. https://doi.org/10.1261/rna.036400.112

Keane, T. M., Goodstadt, L., Danecek, P., White, M. A., Wong, K., Yalcin, B., Heger, A., Agam, A., Slater, G., Goodson, M., Furlotte, N. A., Eskin, E., Nellåker, C., Whitley, H., Cleak, J., Janowitz, D., Hernandez-Pliego, P., Edwards, A., Belgard, T. G., … Adams, D. J. (2011). Mouse genomic variation and its effect on phenotypes and gene regulation. Nature, 477(7364), 289–294. https://doi.org/10.1038/nature10413

Kelleher, E. S., & Barbash, D. A. (2013). Analysis of piRNA-Mediated Silencing of Active TEs in Drosophila melanogaster Suggests Limits on the Evolution of Host Genome Defense. Molecular Biology and Evolution, 30(8), 1816–1829. https://doi.org/10.1093/molbev/mst081

Kim, D., Paggi, J. M., Park, C., Bennett, C., & Salzberg, S. L. (2019). Graph-based genome alignment and genotyping with HISAT2 and HISAT-genotype. Nature Biotechnology, 37(8), 907–915. https://doi.org/10.1038/s41587-019-0201-4

Kneuss, E., Munafò, M., Eastwood, E. L., Deumer, U.-S., Preall, J. B., Hannon, G. J., & Czech, B. (2019). Specialization of the Drosophila nuclear export family protein Nxf3 for piRNA precursor export. Genes & Development, 33(17–18), 1208–1220. https://doi.org/10.1101/gad.328690.119

Krueger, F., & Andrews, S. R. (2016). SNPsplit: Allele-specific splitting of alignments between genomes with known SNP genotypes. F1000Research, 5, 1479. https://doi.org/10.12688/f1000research.9037.2

Langmead, B., Trapnell, C., Pop, M., & Salzberg, S. L. (2009). Ultrafast and memory-efficient alignment of short DNA sequences to the human genome. Genome Biology, 10(3), R25. https://doi.org/10.1186/gb-2009-10-3-r25

Lau, N. C., Seto, A. G., Kim, J., Kuramochi-Miyagawa, S., Nakano, T., Bartel, D. P., & Kingston, R. E. (2006). Characterization of the piRNA complex from rat testes. Science, 313(5785). https://doi.org/10.1126/science.1130164

Li, X. Z., Roy, C. K., Dong, X., Bolcun-Filas, E., Wang, J., Han, B. W., Xu, J., Moore, M. J., Schimenti, J. C., Weng, Z., & Zamore, P. D. (2013). An Ancient Transcription Factor Initiates the Burst of piRNA Production during Early Meiosis in Mouse Testes. Molecular Cell, 50(1). https://doi.org/10.1016/j.molcel.2013.02.016

Liao, H.-F., Kuo, J., Lin, H.-H., & Lin, S.-P. (2016). Isolation of THY1+ Undifferentiated Spermatogonia from Mouse Postnatal Testes Using Magnetic-activated Cell Sorting (MACS). BIO-PROTOCOL, 6(24). https://doi.org/10.21769/BioProtoc.2072

Liao, Y., Smyth, G. K., & Shi, W. (2014). featureCounts: an efficient general purpose program for assigning sequence reads to genomic features. Bioinformatics (Oxford, England), 30(7), 923–930. https://doi.org/10.1093/bioinformatics/btt656

Love, M. I., Huber, W., & Anders, S. (2014). Moderated estimation of fold change and dispersion for RNA-seq data with DESeq2. Genome Biology, 15(12), 550. https://doi.org/10.1186/s13059-014-0550-8

Mohn, F., Sienski, G., Handler, D., & Brennecke, J. (2014). The rhino-deadlock-cutoff complex licenses noncanonical transcription of dual-strand piRNA clusters in Drosophila. Cell, 157(6), 1364–1379. https://doi.org/10.1016/j.cell.2014.04.031

Nellåker, C., Keane, T. M., Yalcin, B., Wong, K., Agam, A., Belgard, T. G., Flint, J., Adams, D. J., Frankel, W. N., & Ponting, C. P. (2012). The genomic landscape shaped by selection on transposable elements across 18 mouse strains. Genome Biology, 13(6), R45. https://doi.org/10.1186/gb-2012-13-6-r45

Ozata, D. M., Gainetdinov, I., Zoch, A., O’Carroll, D., & Zamore, P. D. (2019). PIWI-interacting RNAs: small RNAs with big functions. Nature Reviews Genetics, 20(2), 89–108. https://doi.org/10.1038/s41576-018-0073-3

Özata, D. M., Yu, T., Mou, H., Gainetdinov, I., Colpan, C., Cecchini, K., Kaymaz, Y., Wu, P.-H., Fan, K., Kucukural, A., Weng, Z., & Zamore, P. D. (2020). Evolutionarily conserved pachytene piRNA loci are highly divergent among modern humans. Nature Ecology & Evolution, 4(1). https://doi.org/10.1038/s41559-019-1065-1

Rosenkranz, D., & Zischler, H. (2012). proTRAC--a software for probabilistic piRNA cluster detection, visualization and analysis. BMC Bioinformatics, 13, 5. https://doi.org/10.1186/1471-2105-13-5

Shpiz, S., Ryazansky, S., Olovnikov, I., Abramov, Y., & Kalmykova, A. (2014). Euchromatic Transposon Insertions Trigger Production of Novel Pi- and Endo-siRNAs at the Target Sites in the Drosophila Germline. PLoS Genetics, 10(2), e1004138. https://doi.org/10.1371/journal.pgen.1004138

Siomi, M. C., Sato, K., Pezic, D., & Aravin, A. A. (2011). PIWI-interacting small RNAs: the vanguard of genome defence. Nature Reviews Molecular Cell Biology, 12(4). https://doi.org/10.1038/nrm3089

Song, J., Liu, J., Schnakenberg, S. L., Ha, H., Xing, J., & Chen, K. C. (2014). Variation in piRNA and transposable element content in strains of Drosophila melanogaster. Genome Biology and Evolution, 6(10), 2786–2798. https://doi.org/10.1093/gbe/evu217

Yu, T., Koppetsch, B. S., Pagliarani, S., Johnston, S., Silverstein, N. J., Luban, J., Chappell, K., Weng, Z., & Theurkauf, W. E. (2019). The piRNA Response to Retroviral Invasion of the Koala Genome. Cell, 179(3), 632–643.e12. https://doi.org/10.1016/j.cell.2019.09.002

