## Supplementary Figures for "Genetic polymorphisms lead to major, locus-specific, variation in piRNA production in mouse"

### Supplementary Figure 1

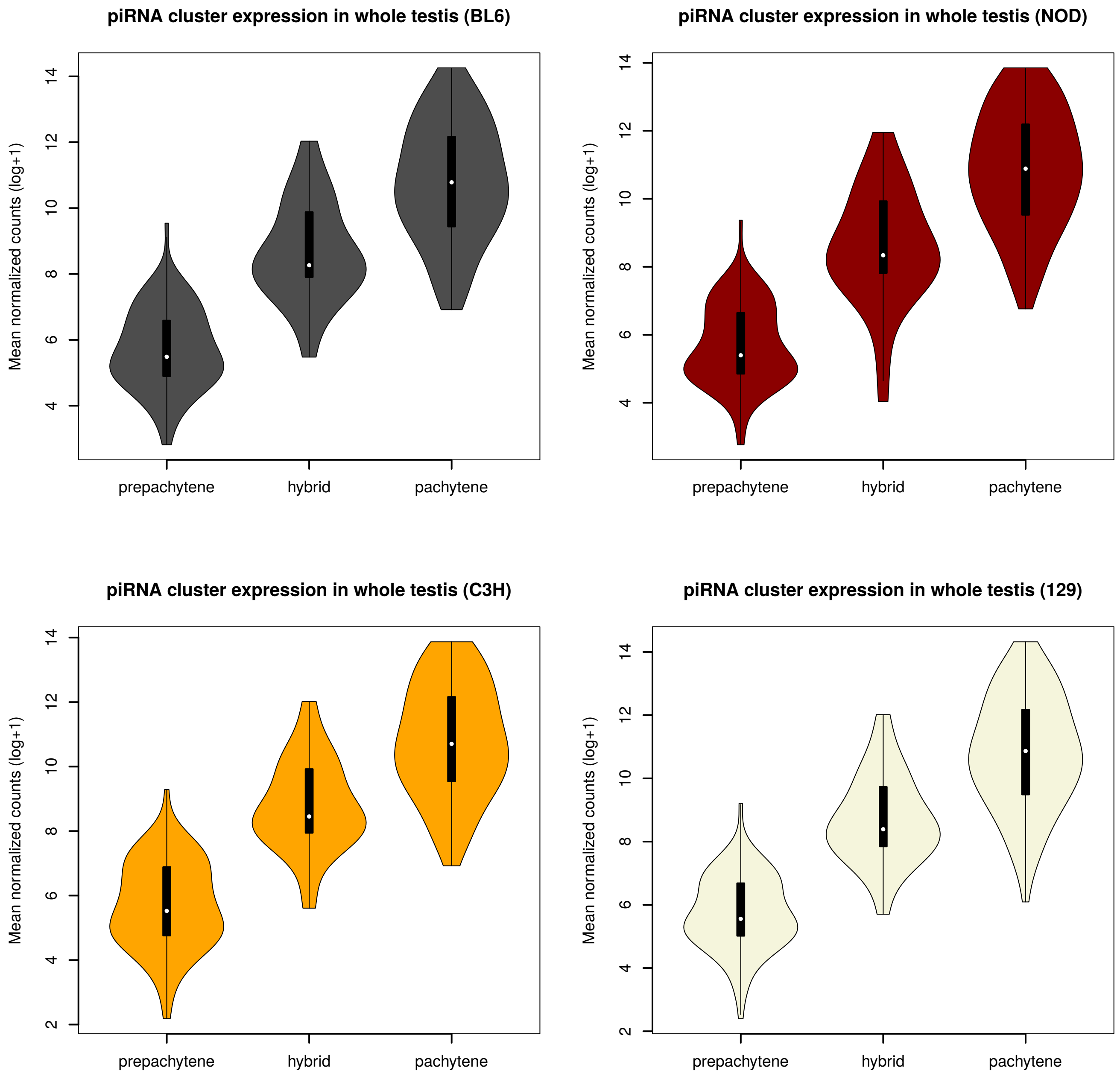

### Supplementary Figure 2A

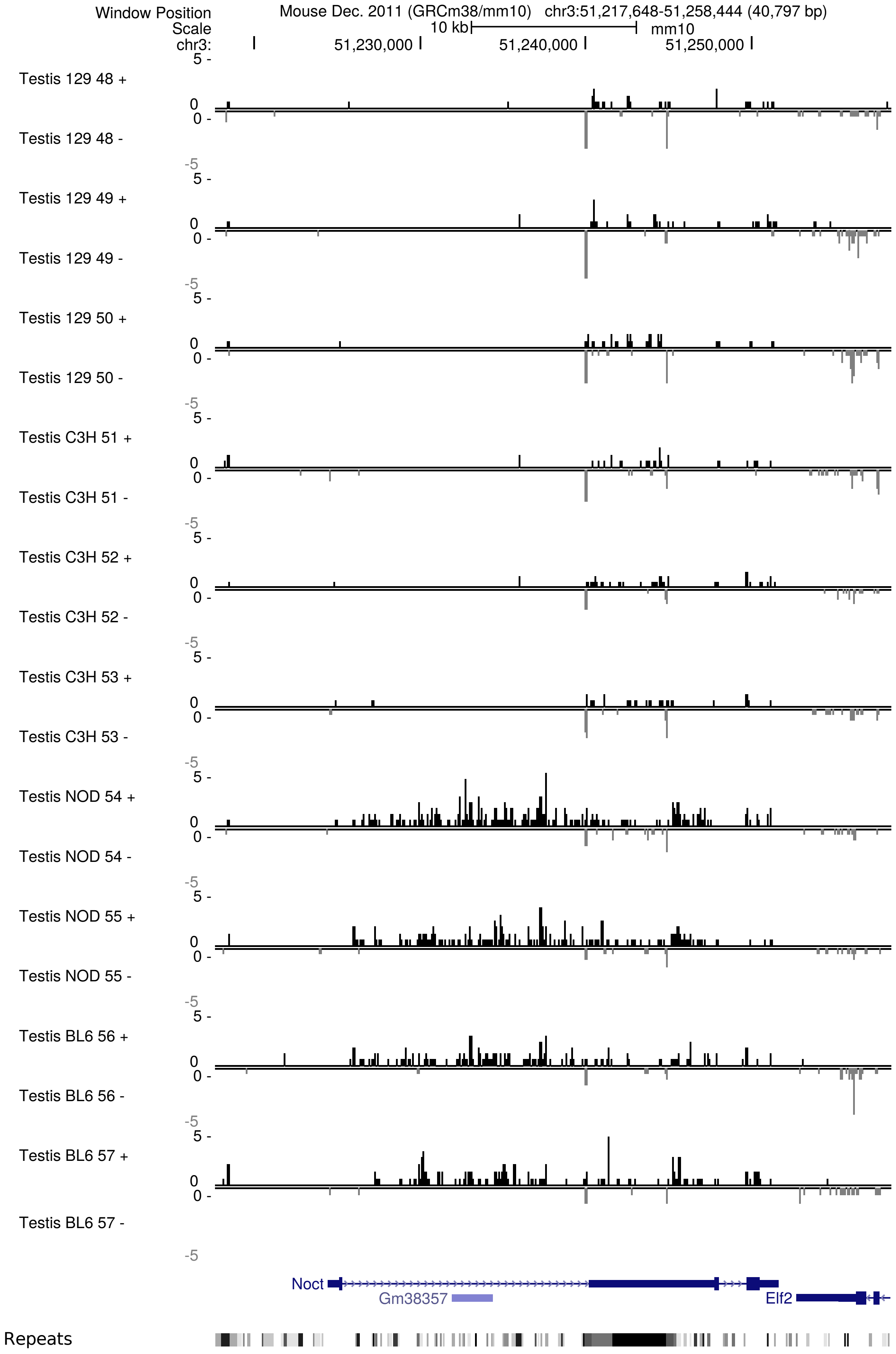

### Supplementary Figure 2B

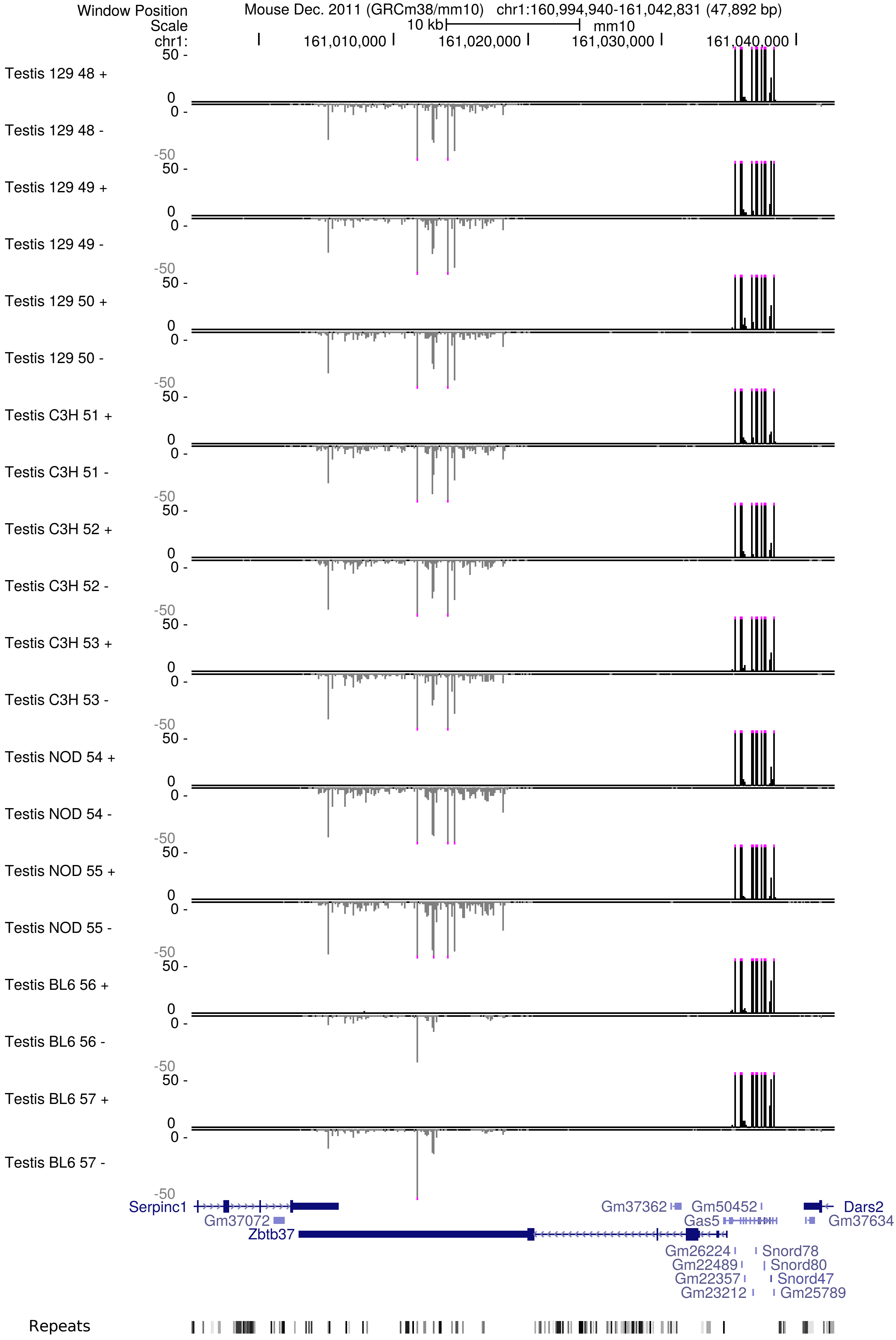

### Supplementary Figure 2C

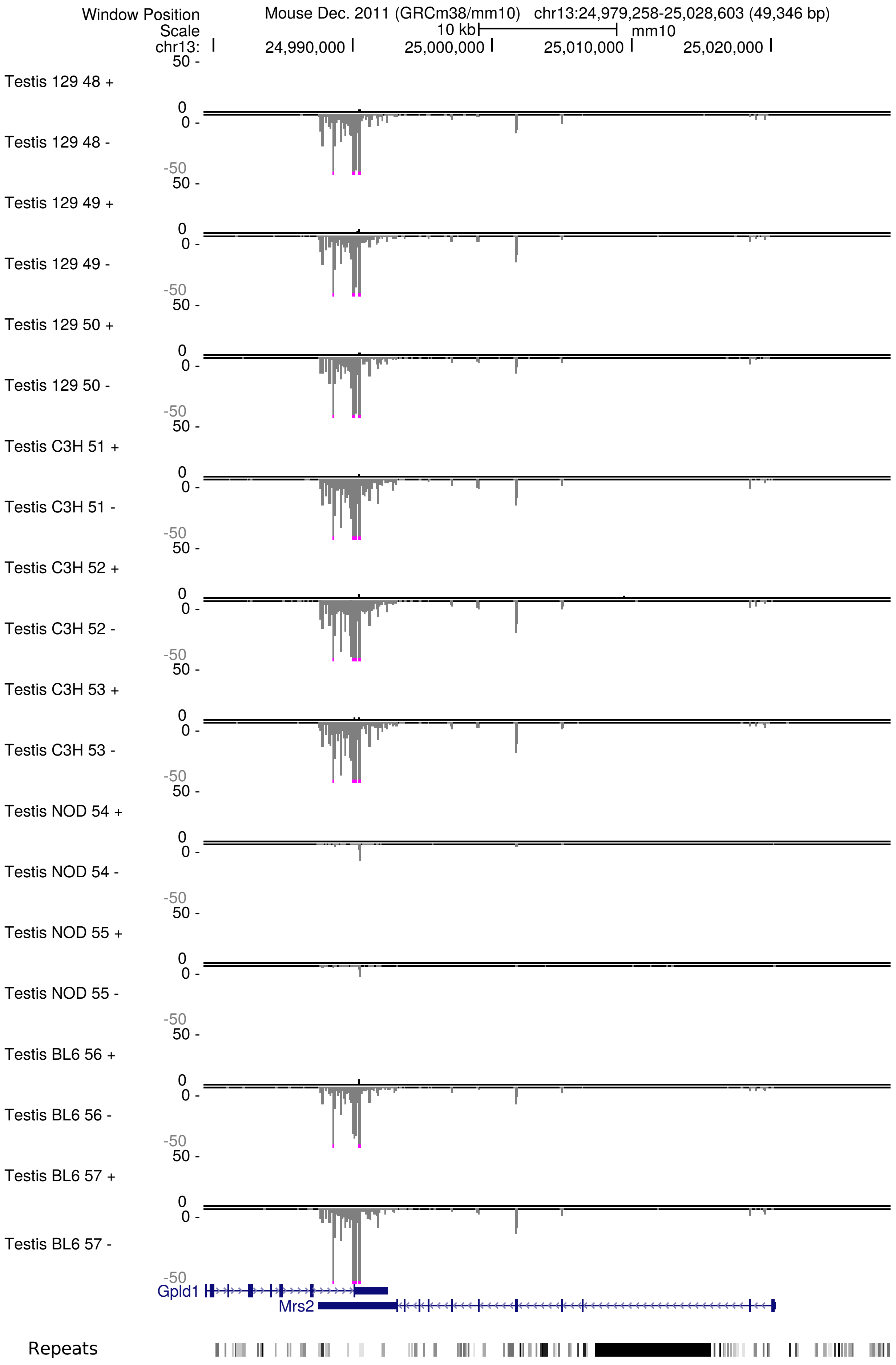

### Supplementary Figure 2D

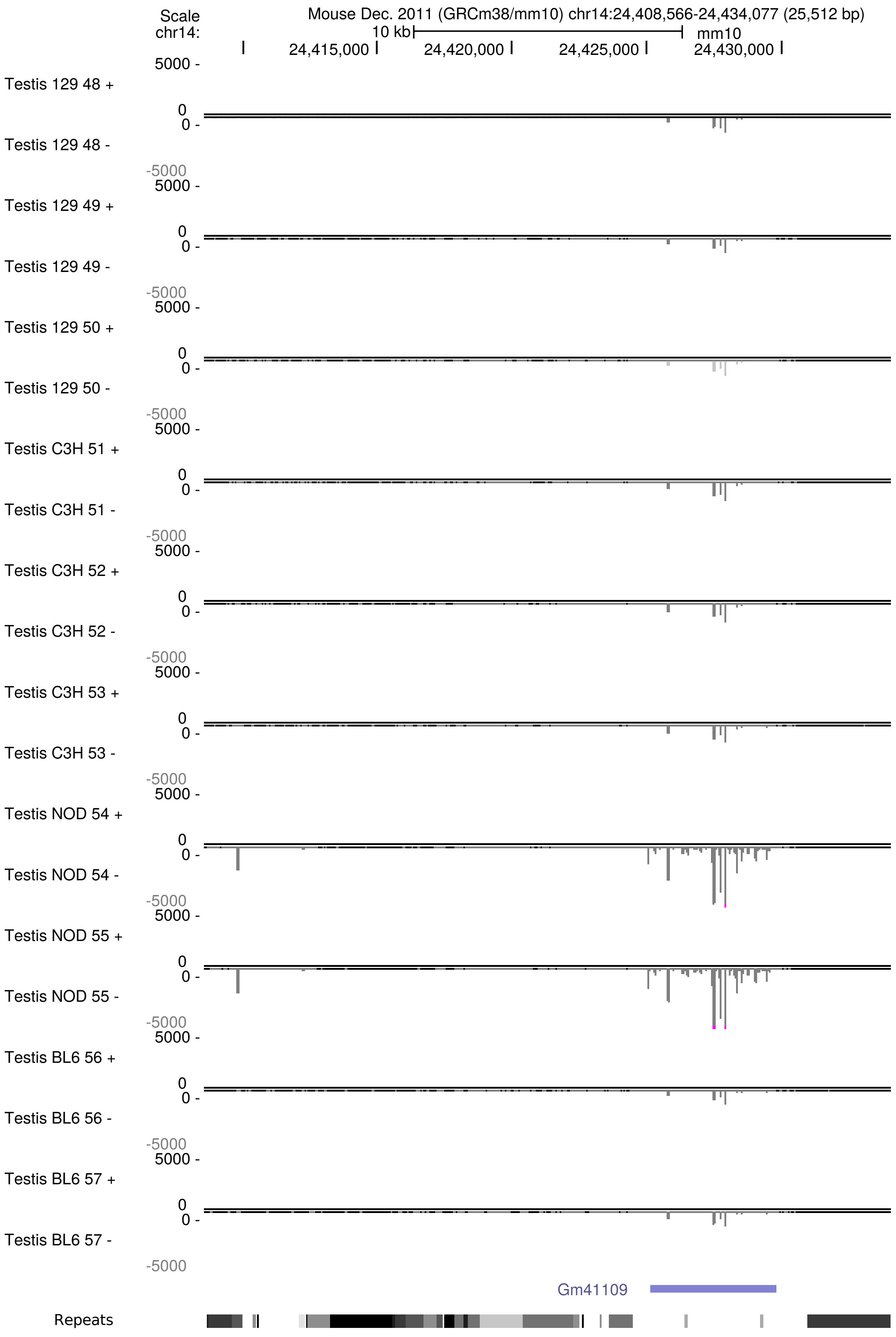

### Supplementary Figure 2E

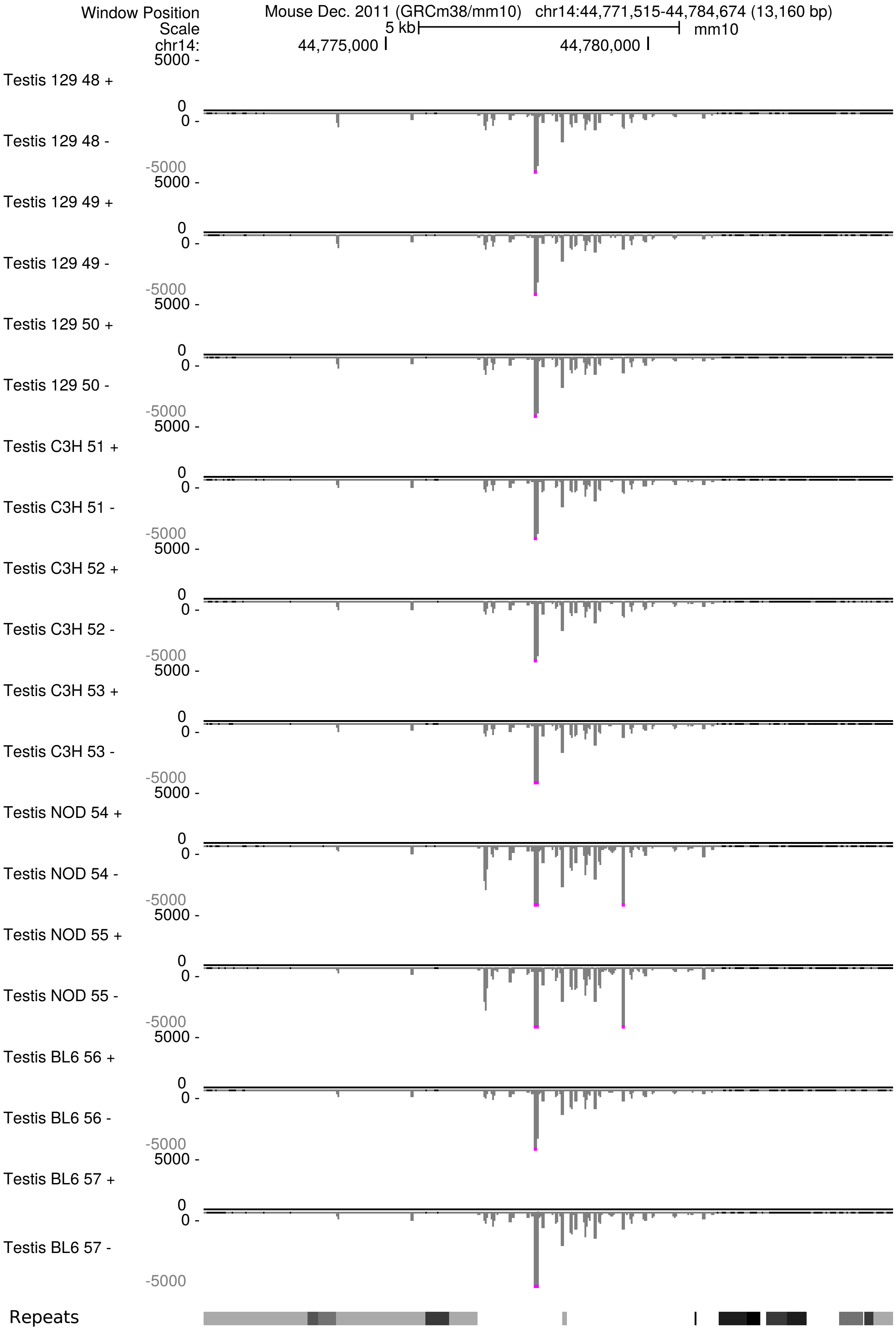

### Supplementary Figure 3

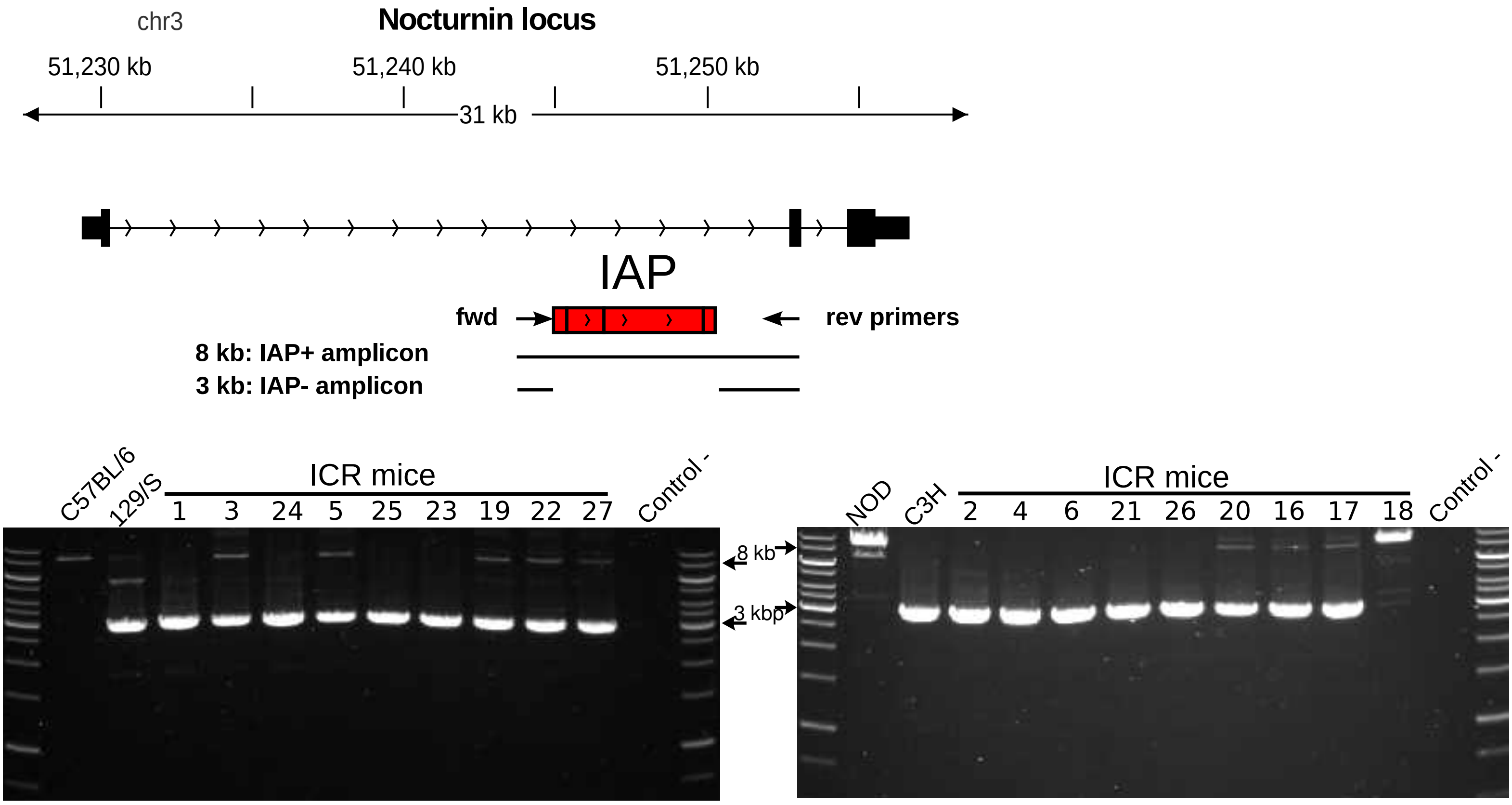
